# Alzheimer’s disease clinical variants show distinct regional patterns of neurofibrillary tangle accumulation

**DOI:** 10.1101/538496

**Authors:** Cathrine Petersen, Amber L. Nolan, Elisa de Paula França Resende, Zachary Miller, Alexander J. Ehrenberg, Maria Luisa Gorno-Tempini, Howard J. Rosen, Joel H. Kramer, Salvatore Spina, Gil D. Rabinovici, Bruce L. Miller, William W. Seeley, Helmut Heinsen, Lea T. Grinberg

**Affiliations:** Memory and Aging Center, Weill Institute for Neurosciences, University of California, San Francisco; Global Brain Health Institute based at University of California, San Francisco and Trinity College, Dublin; LIM-44, University of Sao Paulo Medical School, Sao Paulo, Brazil; Clinic of Psychiatry, University of Würzburg, Wurzburg, Germany; LIM-22, Department of Pathology, University of Sao Paulo Medical School, Sao Paulo, Brazil

**Keywords:** Alzheimer’s disease, neurofibrillary tangles, atypical Alzheimer’s disease, tau pathology, autopsy, human

## Abstract

The clinical spectrum of Alzheimer’s disease (AD) extends well beyond the classic amnestic-predominant syndrome. Previous studies have suggested differential neurofibrillary tangle (NFT) burden between amnestic and logopenic primary progressive aphasia presentations of AD. In this study, we explored the regional distribution of NFT pathology and its relationship to AD presentation across five different clinical syndromes. We assessed NFT density throughout six selected neocortical and hippocampal regions using thioflavin-S fluorescent microscopy in a well-characterized clinicopathological cohort of pure AD cases enriched for atypical clinical presentations. Subjects underwent apolipoprotein E genotyping and neuropsychological testing. Main cognitive domains (executive, visuospatial, language, and memory function) were assessed using an established composite z-score. Our results showed that NFT regional burden aligns with the clinical presentation and region-specific cognitive scores. Cortical, but not hippocampal, NFT burden was higher among atypical clinical variants relative to the amnestic syndrome. In analyses of specific clinical variants, logopenic primary progressive aphasia showed higher NFT density in the superior temporal gyrus (p = 0.0091), and corticobasal syndrome showed higher NFT density in the primary motor cortex (p = 0.0205) relative to the amnestic syndrome. Higher NFT burden in the angular gyrus and CA1 sector of the hippocampus were independently associated with worsening visuospatial dysfunction. In addition, unbiased hierarchical clustering based on regional NFT densities identified three groups characterized by a low overall NFT burden, high overall burden, and cortical-predominant burden, respectively, which were found to differ in sex ratio, age, disease duration, and clinical presentation. In comparison, the typical, hippocampal sparing, and limbic predominant subtypes derived from a previously proposed algorithm did not reproduce the same degree of clinical relevance in this sample. Overall, our results suggest domain-specific functional consequences of regional NFT accumulation. Mapping these consequences presents an opportunity to increase understanding of the neuropathological framework underlying atypical clinical manifestations.

## INTRODUCTION

Pathologically proven Alzheimer's disease (AD) can manifest clinically with a broad spectrum of cognitive presentations beyond the classic progressive amnestic-predominant syndrome. Independent clinicopathological studies have identified AD cases with a constellation of unusual presenting symptoms, generally called atypical or focal cortical AD presentations [17, 33, 64]. These atypical clinical manifestations include corticobasal syndrome (CBS) [23, 35], logopenic variant primary progressive aphasia (lvPPA) [1, 21, 60], posterior cortical atrophy syndrome (PCA) [10, 11], as well as a dysexecutive syndrome resembling behavioral variant frontotemporal dementia (bvFTD) [46, 56]. Atypical clinical variants of AD tend to show a younger age of onset and a lower association with the apolipoprotein E (APOE) ɛ4 allele genotype when compared to typical amnestic cases [16, 45].

The underlying causes of atypical AD presentation remain elusive, but some studies have suggested that co-occurring neurodegenerative pathologies may contribute to the variability in clinical phenotypes of AD cases. For instance, the co-occurrence of Lewy body pathology has been suggested to accelerate cognitive decline and clinical heterogeneity [9, 33]. In the absence of co-pathologies, however, phenotypic variability in pure AD may be a consequence of distinctive patterns of selective neuronal vulnerability at the regional or cellular level [39].

Cognitive decline in AD correlates best with neuronal loss, followed by NFT burden, but only poorly with β-amyloid (Aβ) plaque burden [2, 4, 20, 62]. Atypical clinical variants of AD tend to show more severe cortical atrophy, particularly in key brain areas associated with the most conspicuous clinical feature (e.g. the superior temporal gyrus in lvPPA) [43, 55]. Converging data from quantitative imaging and cerebrospinal fluid studies suggests little difference in the pattern and burden of Aβ pathology distribution between typical and atypical AD presentations [55, 57]. However, studies with tau tracers have found that the level of regional tau burden mirrors the differential pattern of atrophy seen in distinct AD presentations [49, 50, 52, 53]. *In vivo* neuroimaging studies using the tau positron emission tomography tracer ^18^F-AV1451 have shown relevant regional differences in tau uptake among clinical variants of AD. Ossenkoppele et al. reported that individuals with PCA exhibited outsized ^18^F-AV1451 patterns specific to the clinically relevant posterior brain regions, and three out of five individuals with lvPPA showed asymmetric higher left hemisphere ^18^F-AV1451 retention. In addition, regionally relevant ^18^F-AV1451 uptake was associated with domain-specific neuropsychological tests in memory (medial temporal lobes), visuospatial function (occipital, right temporoparietal cortex), and language (left temporoparietal cortex) [52].

Little is known about these differences at the neuropathological and cellular level. Murray et al. advanced the field by describing three distinct AD pathological subtypes based largely on the ratio between hippocampal and cortical NFT density [45]. Individuals with a hippocampal sparing subtype of AD (HpSp) were found to be younger and predominantly male, whereas individuals with a limbic predominant subtype of AD (LP) were older, with a higher proportion of women. HpSp cases were found to include a significantly higher proportion of atypical clinical syndromes; however, it is unclear to what extent this pattern of tau accumulation reflects the regional burden of each specific atypical AD clinical variant [28, 45]. Studies focused on aphasic presentations of AD pathology have noted leftward asymmetry in the cortical NFT burden in aphasic presentations of AD pathology relative to amnestic cases [18, 30]. Mapping the regional burden of tau NFT pathology across multiple clinical variants may further our understanding of the pathological underpinnings of clinical variants of AD.

Here we examined a large postmortem clinicopathological sample to investigate the clinical relevance of regional NFT pathology with regard to multiple clinical variants of AD and domain-specific cognitive decline. We interrogated whether: (i) unique regional distributions of NFT pathology would characterize different clinical variants of AD; and (ii) NFT density in different brain regions would correlate with worse performance on cognitive tests of associated domains. Thus, we systematically mapped the average NFT density throughout selected representative neocortical and hippocampal regions in a pathologically proven AD cohort enriched for atypical presentations of AD.

## MATERIALS AND METHODS

### Participants

All participants were recruited from the clinicopathological cohort of the Neurodegenerative Disease Brain Bank (NDBB) which is part of the Memory and Aging Center at the University of California, San Francisco (MAC/UCSF). At the MAC/UCSF, research participants are followed longitudinally. In our cohort, all individuals underwent in-depth clinical assessment at least once. This assessment included neurological history and examination, as well as comprehensive neuropsychological and functional testing including the Clinical Dementia Rating (CDR). At the NDBB/UCSF, all these participants undergo an extensive dementia-oriented postmortem assessment covering dementia-related regions of interest on the left hemisphere unless upon gross pathology the right was noted to be more atrophic. Neuropathological diagnosis followed currently accepted guidelines [6, 13, 24, 40, 44]. Subtyping for FTLD-TDP and FTLD-tau followed the current “harmonized” nomenclature [37, 38].

From 2005 to 2017, 204 participants who underwent autopsy at UCSF/NDBB received a primary diagnosis of AD pathological changes. From those, we excluded cases with overlapping FTLD (FUS, tau, or TDP-43), chronic traumatic encephalopathy, alpha-synuclein pathology staged Braak ≥ 3, hippocampal sclerosis, or contributing cerebrovascular lesions. Next, to increase the power of the sample, we also included any “pure” AD case with an atypical presentation procured from 2017 to 2018. The final number was 94 cases. The neuropathological investigation was performed blinded to the clinical diagnosis and demographics.

### Clinical history

The clinical syndrome closest to death was determined by chart review and based on published criteria for amnestic syndrome [14, 41, 42], CBS [23, 35], lvPPA [1, 21, 60], PCA [10, 11], and bvFTD [46, 56]. All the charts were reviewed by a behavioral neurologist (ER) blinded to the neuropathological status. If any discrepancies were noted in the diagnosis at different points in time or in different evaluations in the chart, or if there were any atypical clinical features, the chart was reviewed by a second expert behavioral neurologist (ZM) and the final diagnosis was determined after consensus. All participants fit in one of these syndromes, except one who presented with rapid cognitive decline, hyper-somnolence, parkinsonism and ataxia and was classified as dementia without other specification. This participant was excluded from statistical group comparisons of diagnoses. We used the CDR score obtained postmortem with an informant to reflect the participant’s cognition status by death. A diagnosis of very mild dementia required a CDR score of 0.5, and all cognitively normal participants had a CDR = 0 in this evaluation. However, for the analysis, we used the CDR score obtained in the last research visit, aiming to have a reliable picture of the global cognition at the time at which neuropsychological scores were obtained.

We also obtained information for the following variables from the MAC/UCSF Clinical Database and included these in the analysis: age at symptoms onset, age at death, disease duration, sex, and years of education. APOE ɛ4 allele genotyping was done using a TaqMan Allelic Discrimination Assay on an ABI7900HT Fast Real-Time polymerase chain reaction system (Applied Biosystems, Foster City, CA, USA).

We calculated composite z-scores for major cognitive domains, using the neuropsychological evaluation closest to death. The time lag between evaluation and death was corrected for in relevant statistical analyses. This neuropsychological assessment covered four cognitive domains: executive function [design fluency, letter fluency, Stroop test (correct naming), digital backwards, and Trail making B (number of correct lines in one minute)], language ability (Boston Naming Test [32], fluency of animals in one minute, Peabody Picture Vocabulary Test [15], and Information subtest of Verbal IQ from the Wechsler scale), visuospatial ability (modified Rey figure, number location of the Visual Object and Space Perception battery, and block design of the Wechsler Adult Intelligence Scale -III), and memory [California Verbal Learning Test (delayed recall, sum of the learning trials, and recognition accounting for false positives) and modified Rey figure delayed recall]. Performance for each of these four cognitive domains (executive, language, visuospatial, and memory function) was assessed through a predefined calculated composite score averaging the z-scores from the collected neuropsychological raw data. These z-scores are calculated relative to normative data from a cohort of cognitively healthy older adults [61]. These composite scores are used in lieu of specific neuropsychological test performance in order to enhance sensitivity to domain dysfunction and reduce the dimensionality of cognitive assessment data.

### Neuropathological assessment

Using thioflavin-S fluorescent microscopy, quantitative measures of NFT densities were manually assessed with a Zeiss Axio Scan.Z1 fluorescent slide scanner microscope at the Molecular Imaging Center at the University of California, Berkeley. The regions examined for each subject included four neocortical regions: the middle frontal gyrus, superior temporal gyrus, primary motor cortex, and angular gyrus; and two hippocampal regions: the CA1 and subiculum. These regions were chosen as representative association cortices and hippocampal subfields across a range of functional domains and classical vulnerability to AD pathology.

The neuroanatomical sampling design and procedures for microscopy using thioflavin-S fluorescent dye used in this study were informed by techniques developed originally by Terry and colleagues [63]. Briefly, 8μm-thick paraffin-embedded sections were stained using thioflavin-S, and regions of interest were imaged. Three 0.25 mm^2^ areas (500 μm × 500 μm) were sampled at random from each region, and quantitative NFT counts were averaged across these three areas to produce a density score (Supplemental Figure 1). Densities are reported as the number of NFTs per mm^2^.

Thioflavin-S identifies tau NFT pathology as well as β-amyloid neuritic plaques [34]. NFT pathology was distinguished from β-amyloid pathology based on the distinct morphological differences between the aggregates (Figure 1). NFT pathology was distinguished by flame-shaped or globose morphology of fibrous neuronal aggregates. NFT counts included intracellular and extracellular NFTs.

**Fig 1.**
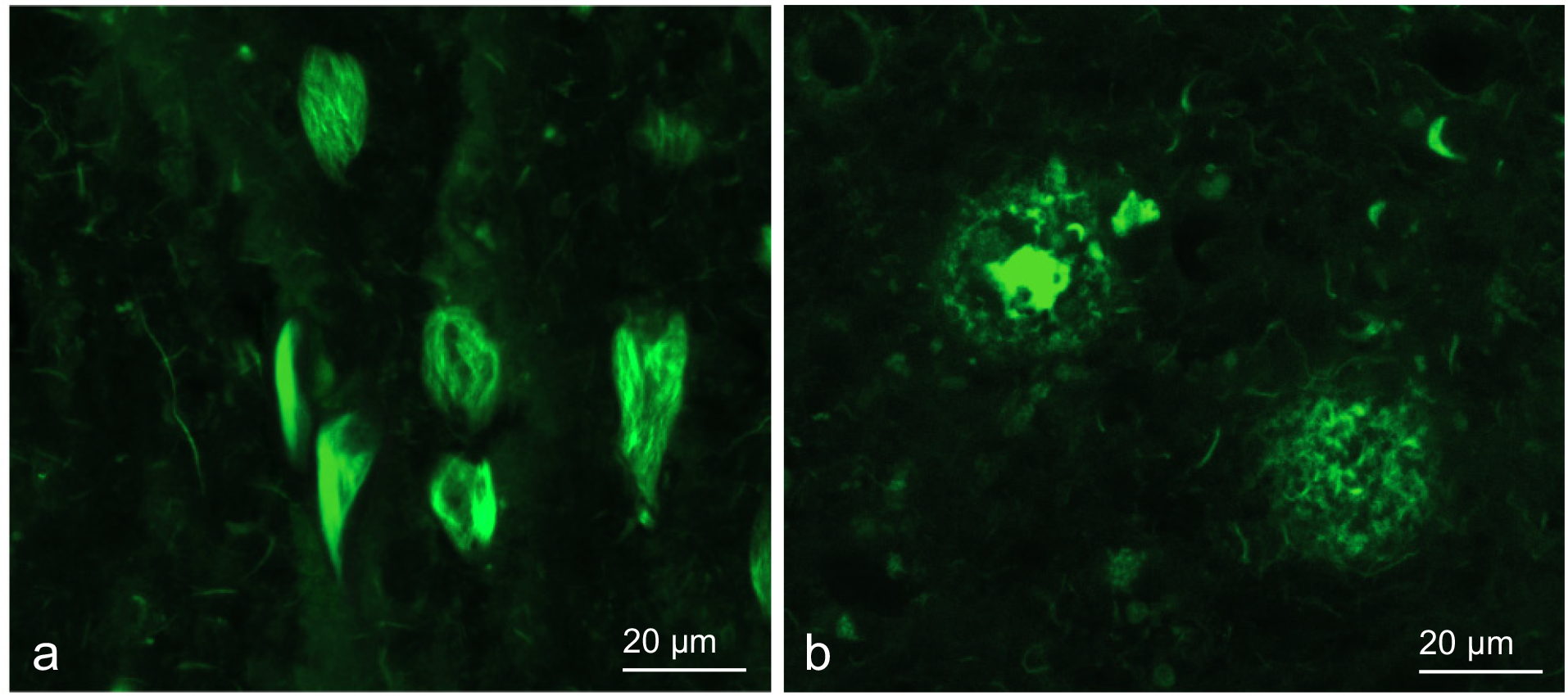
Thioflavin-S identifies tau neurofibrillary tangles (NFTs) and β-amyloid neuritic plaques. We assessed the regional density of NFTs only. **a.** NFT pathology is distinguished by flame-shaped or globose morphology of fibrous neuronal aggregates. **b.** β-amyloid neuritic plaques are distinguished by the processes extending from diffuse or cored plaque aggregates.

### Statistical analysis

To evaluate the differences in cognitive domain composite z-scores and regional NFT densities among the clinical diagnostic groups, we used a Kruskal-Wallis test with *post hoc* pairwise Mann-Whitney U tests. To account for multiple testing, the false discovery rate was set at < 0.05. For these analyses, the cognitively normal/very mild dementia classifications are treated as one diagnostic group. To evaluate the relationships between regional NFT density and demographic covariates, we used multivariate linear regression. Additionally, multivariate linear regression was used to evaluate the relationships between regional NFT density and domain-specific cognitive scores, accounting for clinical covariates and the time lag between neuropsychological evaluation and death.

The above-mentioned supervised statistical learning methods operate best within a hypothesis-driven line of inquiry. While effective, these methods are categorically subject to bias dependent upon the questions asked. In order to analyze the regional NFT density data in an unbiased manner, we supplemented these techniques with unsupervised statistical learning methods. First, to analyze the covariation between NFT density in different brain regions, we used principal component analysis. We abided by Kaiser’s criterion [31], to retain only those factors that have eigenvalues > 1, and Cattell’s criterion [7], which uses a scree plot of eigenvalues and retains all factors in the sharp descent prior to the inflection point. Factor meaningfulness and interpretability were taken into consideration, along with contribution to total variance.

Second, in order to identify patterns of regional NFT accumulation repeated across our sample, we applied an unbiased hierarchical clustering analysis based on the regional NFT density in the hippocampus (CA1 and subiculum) and three association cortices: the middle frontal gyrus, superior temporal gyrus, and angular gyrus. These regions were selected in order to allow effective comparison with existing algorithms for neuropathological subtyping in AD [45]. We validated Ward’s method of hierarchical clustering against other clustering methodologies, such as k-means clustering, based on three internal validation criteria: connectivity, silhouette width, and the Dunn index [12, 19]. The validation was done in R, using the Cluster Validation Package clValid [5].

The results of this unbiased hierarchical clustering analysis were compared to subtypes identified using a previously described threshold-based algorithm. Per Murray et al. [45], subjects were classified as HpSp, LP, and typical AD subtypes based on the density of NFTs in the same regions. The detailed algorithm methods have been previously described. Briefly, to qualify as HpSp, a case must pass three requirements. First, the ratio of the average hippocampal NFT to the average cortical NFT must be less than the 25th percentile of all cases. Second, all of the hippocampal NFT densities must be less than the median values. Third, at least three of the cortical NFT measures must be greater than or equal to the median values. To qualify as LP, a case must pass the converse three requirements. If a case meets criteria for neither HpSp or LP, it is classified as the typical AD subtype. Both the hierarchical clustering and algorithmic subtype analysis used seventy-four cases, all Braak stage > IV to limit the effect of disease progression on group membership and excluding any cases with missing data in the relevant regions.

In order to examine the clinical relevance and applicability of each method of neuropathological partitioning, the resulting groups were contrasted in terms of demographics and neuropsychological composite scores using a Kruskal-Wallis test with *post hoc* Mann-Whitney U-test comparisons. To account for multiple testing, the false discovery rate was again set at < 0.05. A Chi Square test was used to compare the proportion of males, atypical clinical variants, and APOE ɛ4 allele carriers across the groups.

Statistical analyses were performed using R Statistical Software (version 3.4.4; R Foundation for Statistical Computing, Vienna, Austria).

## RESULTS

### Demographics

Of the ninety-four participants, fifty-six (59.6%) were male. The mean (SD) age of symptoms onset was 60.8 (10.5) years, the mean (SD) age of death was 71.2 (10.9) years, and the mean (SD) disease duration was 10.4 (3.7) years. The mean (SD) educational attainment was 15.9 (3.3) years. In 44.2% of the participants, at least one copy of the APOE ε4 allele was present. This cohort predominantly included participants with high AD neuropathologic change [44]. Seventy-six participants (80.9%) were assigned a Braak stage VI for neurofibrillary changes. Eighty-three participants (88.3%) had frequent neuritic plaque pathology by CERAD criteria.

Fifty-two participants (55.3%) were diagnosed with a typical amnestic syndrome, whereas thirty-one (33.0%) were diagnosed with an atypical clinical variant: eight participants were diagnosed with CBS, eight with lvPPA, seven with bvFTD, seven with PCA, and one with an unspecified clinical syndrome. In addition, seven cases met criteria for very mild dementia (CDR = 0.5) and four were cognitively normal at death (CDR = 0). Demographic and clinical characteristics for each diagnostic group are presented in Table 1.

**Table 1.** Demographic characteristics according to clinical diagnostic group

| Clinical diagnosis | n | Proportion male | Mean age of onset (SD) | Mean age of death (SD) | Mean disease duration (SD) | Years of education (SD) | Proportion APOE ε4 carriers |
| --- | --- | --- | --- | --- | --- | --- | --- |
| Amnesic syndrome | 52 | 0.67 | 62.63 (10.53) | 73.21 (10.83) | 10.43 (3.56) | 15.86 (3.91) | 0.54 |
| CBS | 8 | 0.38 | 57.88 (10.05) | 67.88 (9.49) | 10.00 (3.66) | 16.14 (3.08) | 0.25 |
| lvPPA | 8 | 0.38 | 56.13 (5.41) | 66.63 (6.19) | 10.50 (2.00) | 15.63 (2.13) | 0.29 |
| bvFTD | 7 | 0.86 | 51.57 (6.55) | 64.43 (7.18) | 12.86 (5.34) | 15.80 (2.68) | 0.43 |
| PCA | 7 | 0.14 | 53.14 (2.73) | 63.14 (3.80) | 10.00 (3.61) | 15.86 (1.46) | 0.17 |
| Other | 1 | 1.00 | 58.00 ( - ) | 66.00 ( - ) | 8.00 ( - ) | 16.00 ( - ) | 1.00 |
| CN | 7 | 0.43 | 73.43 (7.44) | 82.57 (4.86) | 9.14 (4.81) | 16.43 (2.94) | 0.29 |
| VMD | 4 | 1.00 | - | 86.75 (12.82) | - | 15.25 (1.50) | 0.50 |
| Total | 94 | 0.60 | 60.80 (10.46) | 71.99 (10.87) | 10.43 (3.70) | 15.88 (3.27) | 0.44 |

Abbreviations: APOE, apolipoprotein E; CBS, corticobasal syndrome; lvPPA, logopenic variant primary progressive aphasia; bvFTD, behavioral variant frontotemporal dementia; PCA, posterior cortical atrophy; VMD, very mild dementia; CN, cognitively normal

### Domain-specific cognitive deficits differ among AD clinical variants

Predictably, individuals with lvPPA showed significantly higher impairment on language tasks than individuals diagnosed with a typical amnestic syndrome (p = 0.0021), while individuals with PCA performed significantly worse on visuospatial tasks relative to individuals diagnosed with a typical amnestic syndrome (p = 0.0038). The group of cognitively normal/very mild dementia individuals performed significantly better relative to all other diagnostic groups on executive, language, and visuospatial tasks. For memory tasks, the only significant difference was between the typical amnestic syndrome and the cognitively normal/very mild dementia group (p = 0.0243). Summary statistics are presented in Table 2A.

**Table 2.**
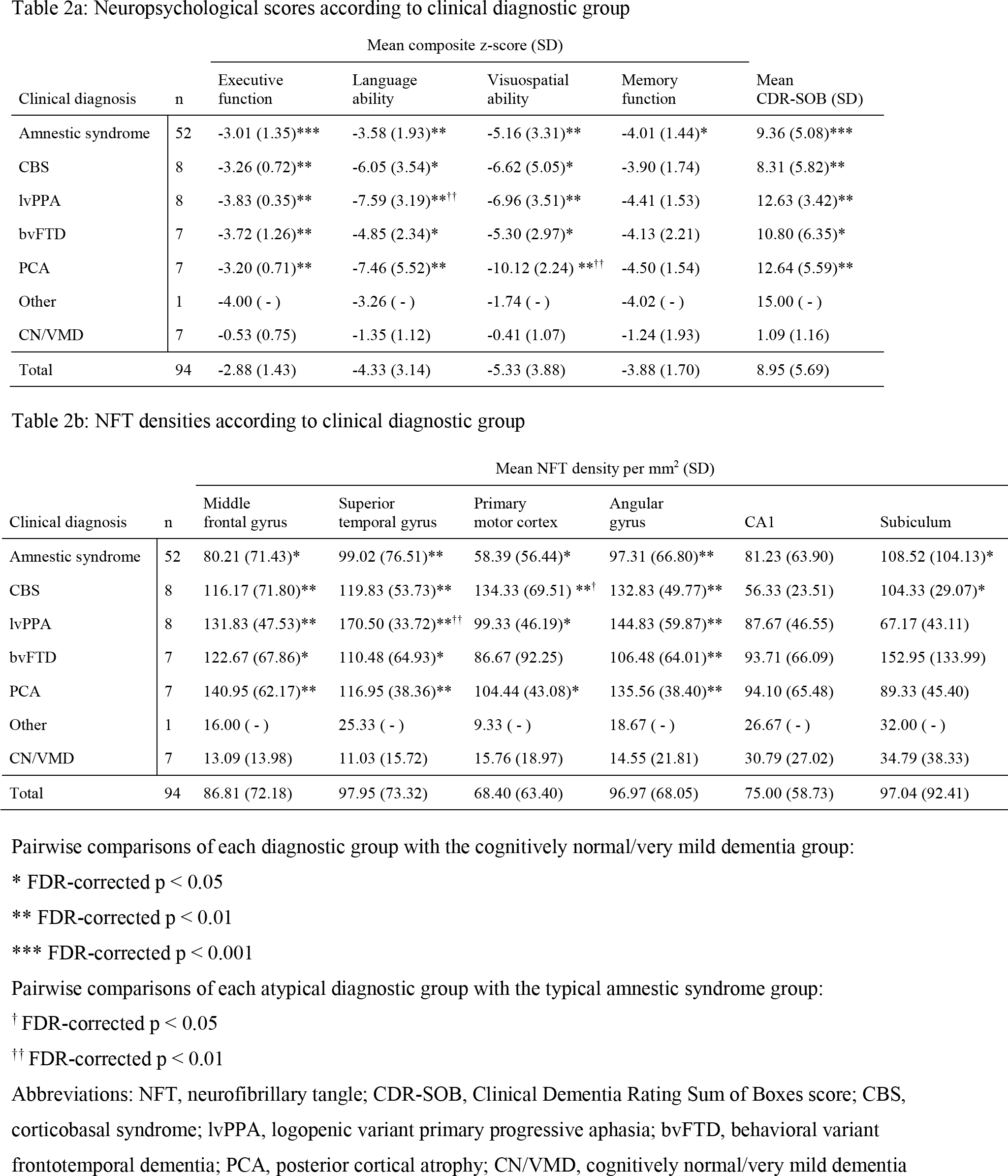
Summary statistics for (**a**) neuropsychological scores and (**b**) regional NFT densities according to diagnostic group. Differences between groups were assessed using a Kruskal-Wallis test with *post hoc* Mann-Whitney U-test pairwise comparisons. The false discovery rate (FDR) was set at < 0.05.

### Distinct regional patterns of NFT accumulation characterize atypical AD clinical variants

The brain regions showing most prominent NFT accumulation differ among AD clinical variants. Figure 2 shows the mean regional NFT density for each diagnostic group. Summary statistics are presented in Table 2B.

**Fig 2.**
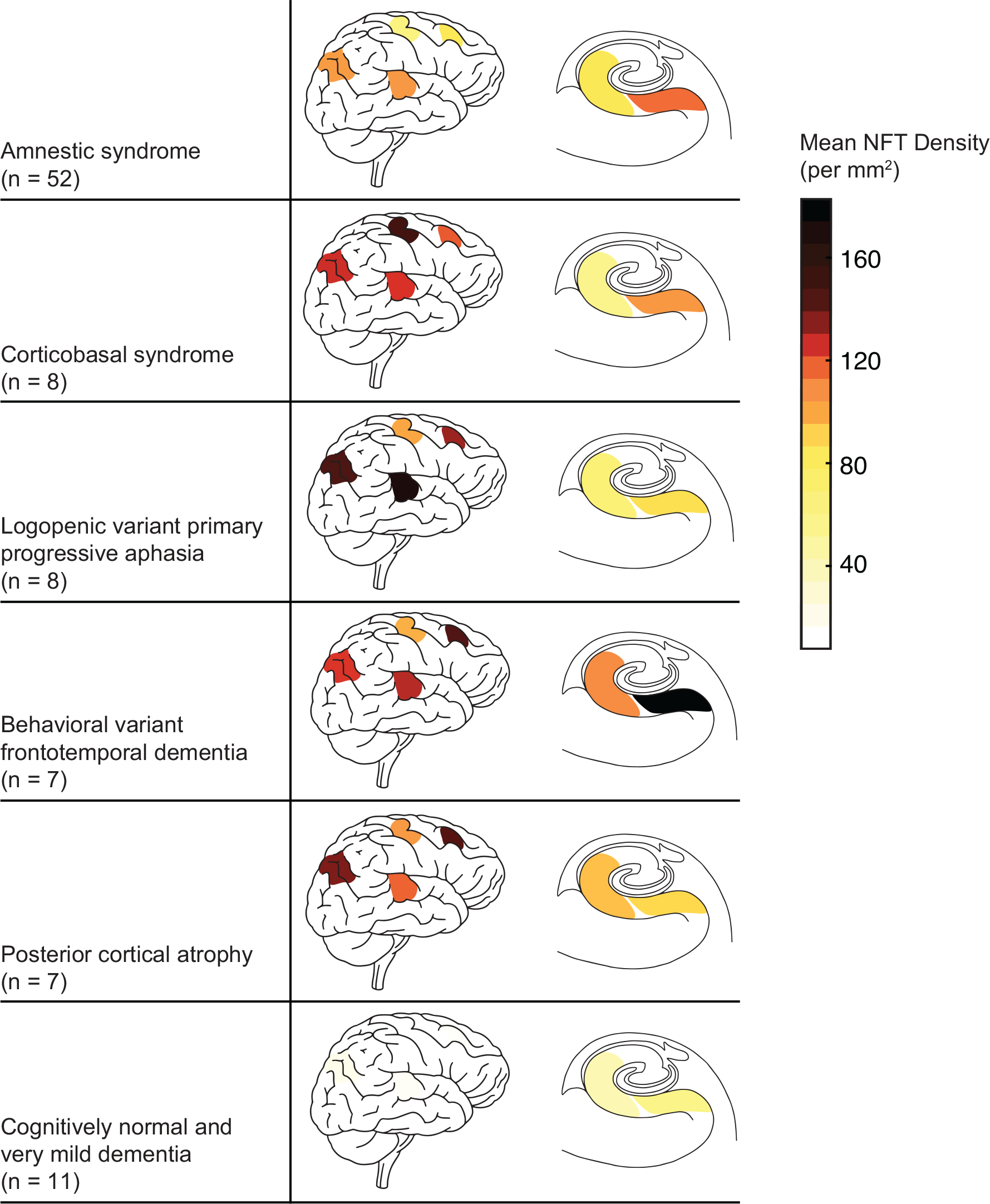
Mean regional NFT densities according to diagnostic group. Regions shown are (left to right): angular gyrus, primary motor cortex (top), superior temporal gyrus (bottom), middle frontal gyrus, CA1, and subiculum. Darker color indicates higher degree of NFT pathology burden. Results of pairwise statistical comparisons are shown in Table 2b.

Notably, individuals with lvPPA showed significantly higher NFT density specific to the superior temporal gyrus relative to individuals with an amnestic syndrome (p = 0.0097) and relative to the other atypical clinical variants (p = 0.0144). Individuals with CBS showed significantly higher NFT density in the primary motor cortex relative to individuals with an amnestic syndrome (p = 0.0205) but not relative to the other atypical clinical variants (p = 0.1544). In our analysis, no other regions were shown to differ significantly between any specific atypical clinical variant and the amnestic syndrome group in this sample; likewise, no other regions differed significantly between any specific atypical clinical variant and the remaining atypical clinical variants. However, groupwise comparisons of the combined atypical clinical variants (n = 31) compared to the amnestic syndrome (n = 52) revealed significantly higher NFT burden in the atypical variant group in each cortical region: the middle frontal gyrus (p = 0.0173), the superior temporal gyrus (p = 0.0452), the primary motor cortex (p = 0.0173), and the angular gyrus (p = 0.0411). In contrast, no significant differences were found for NFT burden in the two hippocampal subfields between the atypical variant group and the typical amnestic syndrome.

In the cognitively normal/very mild dementia group, NFT density was predominantly restricted to the hippocampal subfields. Surprisingly, NFT density in the CA1 subfield of the hippocampus did not significantly differ between any diagnostic groups, whereas NFT density in the subiculum was significantly higher in amnestic syndrome (p = 0.0382) and CBS (p = 0.0336) than in the cognitively normal/very mild dementia group. The cortical regions, however, showed more robust differences between the cognitively normal/very mild dementia group and the remaining diagnostic groups.

### Associations of regional NFT accumulation with neuropsychological performance

Increased cortical NFT burden (an average of the densities in the middle frontal gyrus, superior temporal gyrus, primary motor cortex, and angular gyrus) significantly correlated with more severe cognitive dysfunction for all four domains: executive function (β = −0.0112, p = 0.0033), language ability (β = −0.0287, p = 0.0201), visuospatial ability (β = 0.0276, p = 0.0379), and memory (β = −0.0146, p = 0.0055), correcting for demographic covariates and the time elapsed between the neuropsychological tests and death. In contrast, increased hippocampal NFT burden (an average of the densities in the CA1 and subiculum subfields of the hippocampus) correlated with more severe cognitive dysfunction in just two domains: executive function (β = −0.0054, p = 0.0470) and memory (β = −0.0095, p = 0.0053).

The collinearity of NFT density among the five assessed regions makes it difficult to parse region-specific effects on relevant cognitive domains. Even so, a strong regionally specific effect was observed for visuospatial ability; higher NFT density in the angular gyrus (β = −0.0230, p = 0.0099) and, independently, in the CA1 sector of the hippocampus (β = −0.0184, p = 0.0380) was significantly associated with more severe dysfunction as measured by the visuospatial domain composite z-score, albeit significantly modulated by age of death.

### Cortical and hippocampal axes of variation in tau pathology are clinically relevant

Principal component analysis of NFT regional density data retained two components, together accounting for 78.22% of the variance in regional NFT density. In the resulting plot of variable factor loadings, the four neocortical regions lie close together and the two hippocampal regions lie close together, revealing co-localization of NFT accumulation along two axes: cortical and hippocampal (Supplemental Figure 2A). This dimensionality reduction appears to have some usefulness in discriminating certain atypical clinical variants – namely CBS, lvPPA, and PCA – from the typical amnestic syndrome and cognitively normal/very mild dementia groups, whereas bvFTD is not clearly discriminable (Supplemental Figure 2B).

Multivariate linear regression correcting for demographic covariates sex, disease duration, age of death, years of education, and APOE ε4 allele presence showed a significant inverse correlation of cortical NFT burden with age of death (β = −3.5255, p < 0.0001). Hippocampal NFT burden showed a significant positive association with disease duration (β = 7.5788, p = 0.0002). NFT burden was found to be significantly higher among women in both cortical regions and hippocampal regions (respectively, β = 26.9491, p = 0.0096; β = 35.1174, p = 0.0172), correcting for covariates. There was no significant difference between the regional NFT burden of APOE ε4 allele carriers and non-carriers.

Using unbiased hierarchical clustering, we identified three discrete groups of individuals with varying regional NFT burden (Figure 3A). Clinical and demographic data was not included in the clustering algorithm, which was based solely on regional NFT pathology densities. The resulting groups appear to be characterized by low overall NFT burden (n = 18), high overall burden (n = 21), and cortical-predominant burden (n = 35), respectively. These results are compared to the algorithmic partition into HpSp (n = 5), LP (n = 6), and typical (n = 63) subtypes (Figure 3B). Both the hierarchical clustering and algorithmic partitioning was limited to cases with Braak stage > IV to limit the effect of disease progression on group membership [45]. The clinical associations of the unbiased hierarchical clusters and algorithmic subtypes are summarized in Table 3. HpSp, LP, and typical AD subtypes per Murray et al. differed significantly only along disease duration. Conversely, among the groups identified using hierarchical clustering, we observed significant differences in the frequency of atypical AD clinical variants, sex ratio, age at onset and death, disease duration, executive function, language ability, and CDR scores.

**Fig 3.**
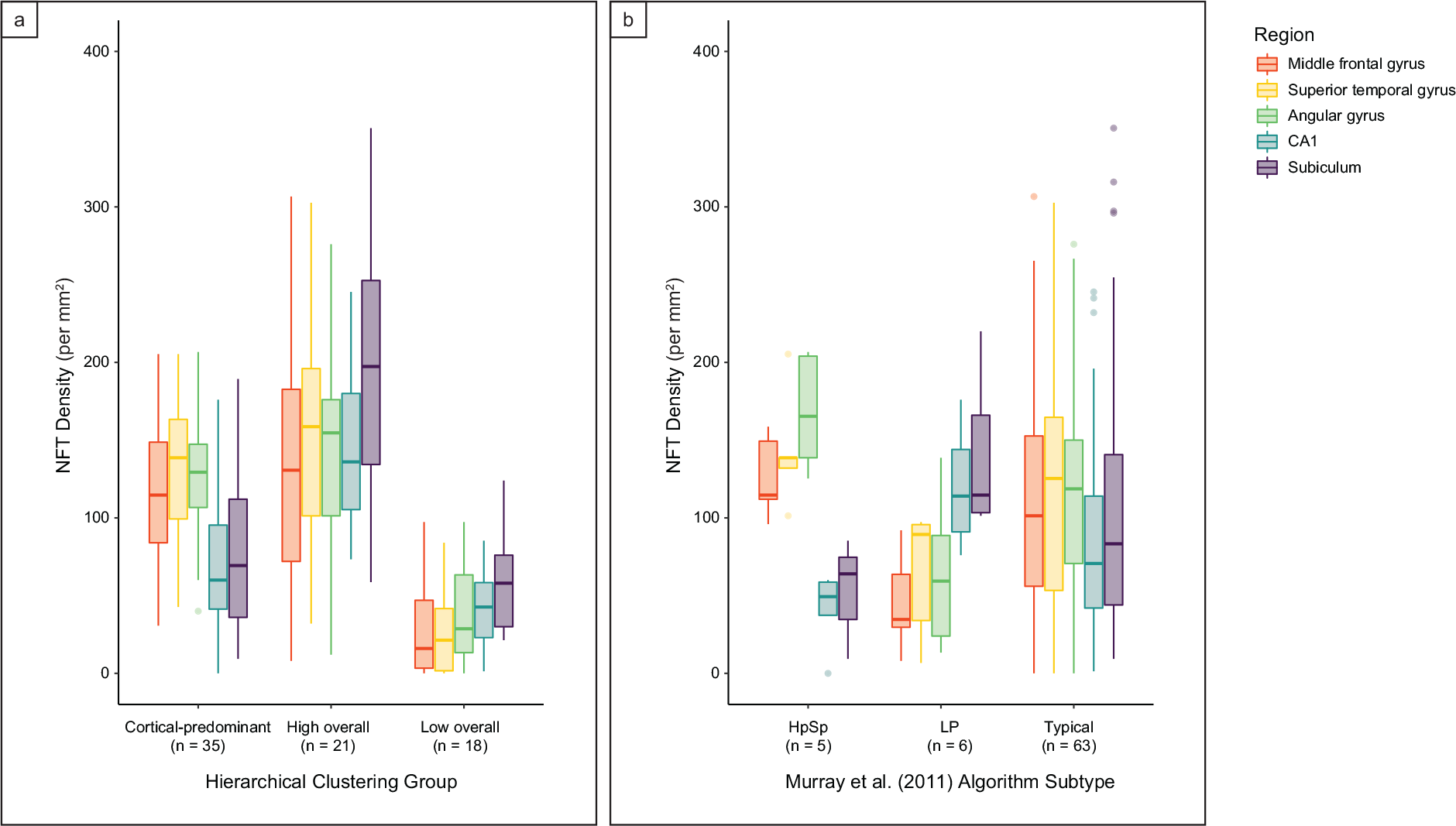
Neuropathological groupings based on regional NFT densities in cases meeting criteria for Braak stage > IV (n = 74). Both methods were based on NFT densities in the regions originally selected by Murray et al. [45]: the middle frontal gyrus, superior temporal gyrus, angular gyrus, CA1, and subiculum. The clinical associations of the hierarchical clusters and algorithmic subtypes are summarized in Table 3. **a.** Unbiased hierarchical clustering identified three clinically relevant clusters of individuals characterized respectively by low overall NFT burden, high overall burden, and cortical-predominant burden. **b.** Typical, hippocampal sparing (HpSp), and limbic predominant (LP) subtypes using a previously defined manual algorithm per Murray et al. [45].

**Table 3.** Clinical and demographic characteristics according to NFT pathological groups defined by unbiased hierarchical clustering and by a previously defined algorithm per Murray et al. [45] in cases meeting criteria for Braak stage > IV (n = 74). Both methods were based on NFT densities in the regions originally selected by Murray et al.: the middle frontal gyrus, superior temporal gyrus, angular gyrus, CA1, and subiculum. Differences between groups were assessed using Kruskal Wallis tests with *post hoc* pairwise Mann-Whitney U-test comparisons for continuous data. Differences for categorial data was assessed using Chi square tests. The false discovery rate (FDR) was set to < 0.05.

| Variable | Hierarchical clustering analysis |  |  |  | Algorithmic subtypes per Murray et al. |  |  |  |
| --- | --- | --- | --- | --- | --- | --- | --- | --- |
|  | Cortical-predominant | High overall | Low overall | *p value* | Hippocampal sparing | Limbic predominant | Typical | *p value* |
| n | 35 | 21 | 18 | - | 5 | 6 | 63 | - |
| Proportion male | 0.46 | 0.43 | 0.94 | *0.0010<sup>b,c</sup>* | 0.60 | 0.67 | 0.56 | *0.8613* |
| Proportion atypical clinical variant | 0.54 | 0.29 | 0.11 | *0.0056<sup>b</sup>* | 0.60 | 0.17 | 0.37 | *0.3312* |
| Proportion APOE ε4 carriers | 0.39 | 0.60 | 0.50 | *0.3399* | 0.20 | 0.83 | 0.47 | *0.0992* |
| Mean age of onset (SD) | 57.69 (8.31) | 55.86 (6.87) | 66.82 (9.36) | *0.0009<sup>b,c</sup>* | 57.60 (6.54) | 62.67 (10.69) | 59.01 (9.21) | *0.5356* |
| Mean age of death (SD) | 66.80 (8.71) | 68.57 (8.08) | 77.22 (10.10) | *0.0020<sup>b,c</sup>* | 65.60 (6.99) | 71.83 (11.02) | 69.98 (9.87) | *0.4074* |
| Mean disease duration (SD) | 9.11 (2.75) | 12.71 (4.17) | 10.41 (3.43) | *0.0070<sup>a</sup>* | 7.00 (2.12) | 9.17 (3.25) | 10.85 (3.65) | *0.0339<sup>d</sup>* |
| Mean years of education (SD) | 15.33 (3.00) | 16.00 (2.13) | 17.06 (3.27) | *0.1884* | 14.00 (4.69) | 16.00 (2.83) | 16.10 (2.73) | *0.7282* |
| Mean CDR-SOB (SD) | 8.03 (4.66) | 12.11 (5.65) | 9.50 (5.85) | *0.0455<sup>a</sup>* | 9.00 (4.30) | 8.25 (3.57) | 9.63 (5.70) | *0.8307* |
| Memory | 1.26 (0.68) | 2.03 (0.95) | 1.47 (0.81) | *0.0171<sup>a</sup>* | 1.30 (0.67) | 1.67 (0.52) | 1.52 (0.88) | *0.6868* |
| Orientation | 1.14 (0.82) | 1.92 (0.95) | 1.28 (0.86) | *0.0146<sup>a</sup>* | 1.40 (1.08) | 1.67 (0.82) | 1.55 (0.89) | *0.6072* |
| Judgment & problem solving | 1.33 (0.66) | 1.92 (1.00) | 1.42 (0.93) | *0.0889* | 1.30 (0.67) | 1.25 (0.61) | 1.55 (0.89) | *0.7260* |
| Community affairs | 1.44 (0.77) | 1.97 (0.98) | 1.42 (0.86) | *0.1055* | 1.60 (0.89) | 1.42 (0.66) | 1.60 (0.90) | *0.9286* |
| Home & hobbies | 1.33 (0.79) | 2.00 (1.04) | 1.67 (0.95) | *0.0628* | 1.30 (0.67) | 1.42 (0.66) | 1.63 (0.98) | *0.7159* |
| Personal care | 0.97 (0.95) | 1.74 (1.23) | 0.78 (1.11) | *0.0186<sup>a,c</sup>* | 1.00 (0.71) | 0.83 (0.75) | 1.16 (1.18) | *0.9051* |
| Mean executive z-score (SD) | -3.13 (0.98) | -3.72 (0.96) | -2.40 (1.70) | *0.0223* | -3.49 (0.92) | -2.29 (0.69) | -3.14 (1.31) | *0.1573* |
| Mean language z-score (SD) | -4.65 (3.28) | -5.17 (2.99) | -3.06 (1.72) | *0.0904* | -3.73 (2.03) | -2.85 (1.04) | -4.54 (3.07) | *0.4453* |
| Mean visuospatial z-score (SD) | -6.16 (3.62) | -6.79 (3.38) | -3.66 (3.77) | *0.0360* | -7.27 (3.25) | -4.85 (3.80) | -5.63 (3.80) | *0.5759* |
| Mean memory z-score (SD) | -3.89 (1.73) | -4.59 (1.10) | -3.55 (1.64) | *0.0780* | -3.98 (0.95) | -4.23 (1.12) | -3.99 (1.66) | *0.8668* |

a - Pairwise comparison of “cortical-predominant” cluster and “high overall” cluster, FDR-corrected $p < 0.05$
b - Pairwise comparison of “cortical-predominant” cluster and “low overall” cluster, FDR-corrected $p < 0.05$
c - Pairwise comparison of “low overall” cluster and “high overall” cluster, FDR-corrected $p < 0.05$
d - Pairwise comparison of typical AD subtype and HpSp AD subtype, FDR-corrected $p < 0.05$
Abbreviations: NFT, neurofibrillary tangle; AD, Alzheimer’s disease; APOE, apolipoprotein E; CDR-SOB, Clinical Dementia Rating Sum of Boxes score

## DISCUSSION

In this study, we aimed to interrogate the impact of regional NFT distribution and burden on the clinical expression of AD by investigating a cohort of pathologically proven AD cases including four different atypical clinical presentations as well as the typical amnestic presentation. In addition, we used the same cases to investigate a possible correlation between NFT burden and cognitive scores, regardless of the presentation syndrome. Our study unveiled the following novel findings: (i) NFT regional burden aligns with the clinical presentation across multiple different syndromes and also with region-specific cognitive scores, (ii) unbiased hierarchical clustering based on regional NFT densities identified groups showing more clinical relevance than previously suggested models for classifying AD cases neuropathologically. Moreover, we observed that the pattern of cortical and hippocampal NFT burden correlates with specific demographics, in line with previous studies.

Among our cases, the regional NFT burden varied considerably. Such variation showed a strong link to the clinical presentation. NFT burden in all four cortical regions examined was found to be significantly higher among atypical clinical variants relative to the typical amnestic syndrome, suggesting that a high cortical burden may be generalizable among atypical clinical variants of AD.

In analyses of specific clinical variants, we observed that lvPPA and CBS may show exacerbated NFT burden in specific cortical regions beyond this collective heightened cortical burden. Individuals with lvPPA, a syndrome clinically characterized by predominant impaired single-word retrieval and impaired repetition with relative sparing of memory functions [1, 21, 22, 60], showed significantly higher NFT density in the superior temporal gyrus relative to individuals diagnosed with an amnestic syndrome. The superior temporal gyrus is implicated in early cortical stages of speech perception [26, 58], and our results show confirmation of elevated tau burden in this region in patients manifesting lvPPA, as has been suggested by tau PET imaging and neuropathological studies comparing aphasic and amnestic AD presentations [18, 30, 49, 52]. While previous studies have focused on the comparison of lvPPA with amnestic AD, our study unveiled that lvPPA also shows significantly higher NFT density in the superior temporal gyrus relative to the other atypical clinical variants. The left hemisphere was evaluated for all cases of lvPPA, and most other cases (64%), but we wanted to avoid any confounding effect of comparison with right hemisphere superior temporal gyrus densities due to the localization of language regions in the left hemisphere. Thus, we repeated both of these comparisons limited to cases in which the left hemisphere was evaluated, and retained significance for both.

Furthermore, individuals with CBS, a syndrome featuring initial asymmetric rigidity and apraxia, extrapyramidal dysfunction and symptoms of pericentral cortical involvement [3, 23, 35], showed significantly higher NFT density in the primary motor cortex relative to individuals diagnosed with an amnestic syndrome. Perhaps due to the limited sample size, this difference was not significant when compared to the remaining atypical clinical variants, and further research is necessary to confirm whether heightened NFT burden in the primary motor cortex is unique to CBS cases. Although our results are not necessarily surprising, our paper unveils neuropathological correlates of the differential regional involvement observed by neuroimaging methods among multiple AD clinical variants and emphasizes the relevance of tau pathology as a determinant of such variations.

Interestingly, average CA1 NFT burden did not reach very high levels in any of the diagnostic groups, and there was not sufficient statistical evidence to distinguish NFT burden in CA1 in each clinical variant from that of the cognitively normal/very mild dementia group. Conversely, NFT density in the subiculum was significantly higher in amnestic syndrome and CBS than in the cognitively normal/very mild dementia group, highlighting the importance of subdividing the hippocampal formation using proper anatomical classification in any kind of research in AD. Of note, although the average NFT burden in the subiculum appears to be very high in the bvFTD group, the relatively small sample size and broad range of NFT density among the cases (one individual had only negligible tau pathology) precluded statistical significance. In any case, the tendency of the subiculum to show higher NFT burden in bvFTD cases is intriguing and warrants further research once a larger sample is available, particularly because some studies suggest a possible role for the subiculum in temporal behavioral control, although this possibility has yet to be fully explored [8, 25, 47]. Nevertheless, few if any fibers directly connect the lateral frontal cortex with the subiculum in macaques [27]. If the same applies to humans, a possible direct tau spread between the two regions would be unlikely.

Previous studies have suggested that relative sparing of the hippocampal formation coupled with exacerbated cortical tau pathology may distinguish atypical clinical variants of AD from the typical amnestic syndrome [45, 54]. These studies implicate the relative burden of hippocampal compared to cortical NFT pathology as an important potential driver of atypical presentations. However, in our study we failed to observe any significant differences in hippocampal NFT density (both CA1 and subiculum) between amnestic and atypical syndromes. Our results suggest that the heightened cortical tau accumulation observed in atypical AD clinical variants may be independent of the hippocampal tau burden. To clarify these results, the next step would be to test these assumptions in an equally large independent sample.

Discrepancies between expected hippocampal and cortical NFT burden lead to the question of whether atypical cases follow the same stereotypical progression proposed by Braak and Braak, which has been reproduced in multiple studies focused on typical AD cases. To this end, investigating cognitively normal/very mild dementia groups could prove very informative. If cases with an atypical presentation fail to follow the Braak scheme, a small number of cases at early stages of AD pathology should show NFTs in specific cortical areas in the absence of NFT accumulation in subcortical regions and the hippocampal formation. This scenario was not observed in any of our cognitively normal/very mild dementia cases. However, we only have eleven cases in this group, and those cases tended to be older and thus far more prone to typical clinical manifestations of AD, making our cohort less than ideal to explore these questions. Studies using population-based clinicopathological cohorts containing relatively young cases would be more appropriate to such investigation. Regardless, this remains an open question in the field.

In addition to comparing NFT regional burden among AD differential clinical phenotypes, we also investigated possible correlations of regional NFT burden with cognitive scores. Notably, higher NFT density in the angular gyrus and CA1 were independently associated with more severe visuospatial dysfunction. These results corroborate the strong association of regional NFT burden with domain-specific functions, as the angular gyrus is implicated in spatial cognition, and the CA1 has more recently been implicated in spatial encoding in addition to its well-documented role in memory [29, 36, 59].

Classically, AD pathology has been considered a homogeneous entity. This assumption was challenged by Murray and colleagues, who created an algorithm based on the thresholds of NFT burden in selected neocortical and hippocampal regions which classified AD in three distinct neuropathological patterns, namely a typical pattern and less frequent HpSP and LP patterns [45]. In their assessment, they found that these subtypes showed differences in terms of gender ratio, age, disease duration, and percentage of atypical presentations. To assess whether distinct patterns of regional NFT density were also present in our cohort, we first applied the same algorithm proposed by Murray et al. to our cases. We succeeded in classifying our cases per Murray at al. algorithm, with a similar proportion of cases per subtype. However, when using this classification scheme the only observed differences among the subtypes were in disease duration; in which the HpSp subtype showed a shorter disease duration than typical subtype. Of note, age of onset, age of death, and the frequency of atypical AD clinical variants appeared to follow the same trend as described by Murray et al., however, differences were not statistically significant.

For comparison, we used the same regions and type of data as the algorithm per Murray et al. to apply unbiased hierarchical clustering analysis. Our unbiased hierarchical clustering analysis also identified three clusters; however, they were notably different from those achieved with the previous algorithm and appeared to be characterized by low overall NFT burden, high overall burden, and cortical-predominant burden, respectively. Interestingly, the hierarchical clustering groups showed high relevance to clinical characteristics. We found significant differences in the frequency of atypical AD clinical variants, gender ratio, age at onset and death, disease duration, executive function, language ability, and CDR scores. The frequency of atypical AD clinical variants was highest in the cortical-predominant group (54%). Also of interest, our groupings based on hierarchical clustering showed significant differences on CDR-sum of boxes scores for memory, orientation and personal care even in pairwise comparisons of the high overall burden and cortical-predominant groups.

We propose that a primary difference between the two methods may be that manually defined algorithms rely on assumptions about what can be considered “typical” and which differences warrant distinction. The algorithm proposed by Murray et al. constructs the thresholds for each classification along expectations of which cases should fit an atypical profile, on the assumption that the ratio between hippocampal and cortical NFT density is an important driver of clinical heterogeneity. In doing so, it achieves highly dichotomized pathologic profiles in each “atypical” group but risks masking the spectrum of pathologic presentation. Thus, the lack of significant results may be partially due to a lack of statistical power, which limits the applicability of this algorithm to other series. In contrast, the strength of our hierarchical clustering method is its unbiased approach. The characteristic patterns within our hierarchical clustering groups implicate specific regional accumulation of NFT as a contributor to the clinical presentation of AD, which may be contingent on differential regional vulnerability to tau pathology. However, our results using hierarchical clustering suggest that clinically relevant pathologic groupings may be more nuanced than previously suggested. Next steps would ideally involve testing our clustering method in other series containing typical and atypical cases. It is possible that hierarchical clustering analysis could produce different results when applied to other series; however, unbiased approaches are consistently most effective in detecting meaningful differences that are generalizable.

In addition to the novel findings discussed above, we observed predicted clinical associations of cortical and hippocampal NFT accumulation. We found significantly lower cortical NFT burden with increasing age, and both cortical and hippocampal NFT burden differed significantly by sex, with women showing a higher NFT burden. Surprisingly, we found no significant difference in regional NFT density between APOE ɛ4 allele carriers and non-carriers, likely due to the high proportion of Braak stage VI cases in this sample. Braak stage VI has been shown to be overrepresented among APOE ɛ4 allele carriers, relative to non-carriers [48]. In our cases, APOE ɛ4 allele carriers appeared to be underrepresented among atypical clinical syndromes; however, these differences were not statistically significant.

We acknowledge that our study has limitations. Our insight into specific atypical clinical syndromes was limited by the number of subjects within each diagnosis. However, all cases were characterized in depth clinically and neuropathologically. The MAC/UCSF specializes in atypical dementia, and the raw neuropsychological test data was comprehensive, allowing for well-informed assessments of cognitive dysfunction based on composite scores. Similar methods for obtaining z-scores for each domain have been extensively used in previous publications [51, 52]. Additionally, although our sample size might have been doubled had we not excluded cases with confounding mixed pathology (e.g. Lewy Body Disease was present in many of the excluded cases), we felt that it was crucial to focus on pathologically pure AD cases in order to more effectively discriminate correlates of regional NFT burden. Furthermore, we chose to focus on six neocortical and hippocampal regions because of their relevance across a range of functional domains and classical vulnerability to AD pathology. Future work to further elucidate the relationship between regionally specific tau accumulation and clinical heterogeneity in AD would benefit from assessing additional brain regions, as well as intra-regional differences such as NFT pathology in layers III and V, or posterior and anterior areas within particular regions such as the subiculum, which have been associated with functionally distinct roles. Finally, clustering analysis is a method under ongoing constant improvement and therefore, there are several proposed methods available, none of which are universally applicable. For this reason, we selected Ward’s method of hierarchical clustering based on a series of validation steps in accordance with three well-documented indices of internal validation, assessing the compactness, connectedness, and separation of the cluster partitions, as detailed in Methods.

In summary, this study highlights the clinical relevance of regional patterns of NFT accumulation in AD cases, suggesting domain-specific functional consequences of regional NFT accumulation. These results expand on past findings from neuroimaging, postmortem, animal, and cerebrospinal fluid studies suggesting that regional NFT aggregation is closely linked to the clinical manifestations of AD. Continued work to map the regionally specific clinical consequences of tau accumulation presents an opportunity to increase understanding of the neuropathological framework underlying atypical clinical manifestations. In particular, the underlying mechanisms connecting NFT pathology to regional selective vulnerability are unknown, and comparing typical and atypical AD presentations may present an ideal framework to explore this question.

## DECLARATIONS

### Compliance with ethical standards

This study was approved by the UCSF Institutional Review Board (reference number) and all the participants or their legal representatives signed a written informed consent that was obtained according to the 1964 Declaration of Helsinki and its further amendments.

### Conflict of interest

The authors have no duality or conflicts of interest to declare.

### Data Availability

The datasets used and analyzed during the current study are available from the corresponding author upon reasonable request. Raw data is provided as supplementary material.

## Acknowledgments

This study was supported by the National Institute of Health grant K24AG053435 and institutional grants P50AG023501, P01AG019724. MLGT was funded by the National Institute of Health grants K24DC015544A and R01NS50915.

The authors thank the patients and their families for their invaluable contribution to brain aging neurodegenerative disease research. ER is an Atlantic Fellow for Equity in Brain Health and thanks the fellowship for supporting her work. This study was supported by the National Institute of Health grant K24AG053435 and institutional grants P50AG023501, P01AG019724. MLGT was funded by the National Institute of Health grants K24DC015544A and R01NS50915.

