## Supplemental figures for "Alzheimer’s disease clinical variants show distinct regional patterns of neurofibrillary tangle accumulation"

**Journal:** *Acta Neuropathologica*

**Send correspondence to:**

Lea Tenenholz Grinberg, MD, PhD, Associate Professor in Residence

Memory and Aging Center, Weill Institute for Neurosciences, University of California, San Francisco

**Supplemental Fig 1** Example of the neuropathological sampling scheme from the subiculum in one individual. Three 0.25 $mm^2$ areas (500 μm x 500 μm) are sampled at random, and quantitative NFT counts were averaged across these three areas to produce a density score reported per $mm^2$.

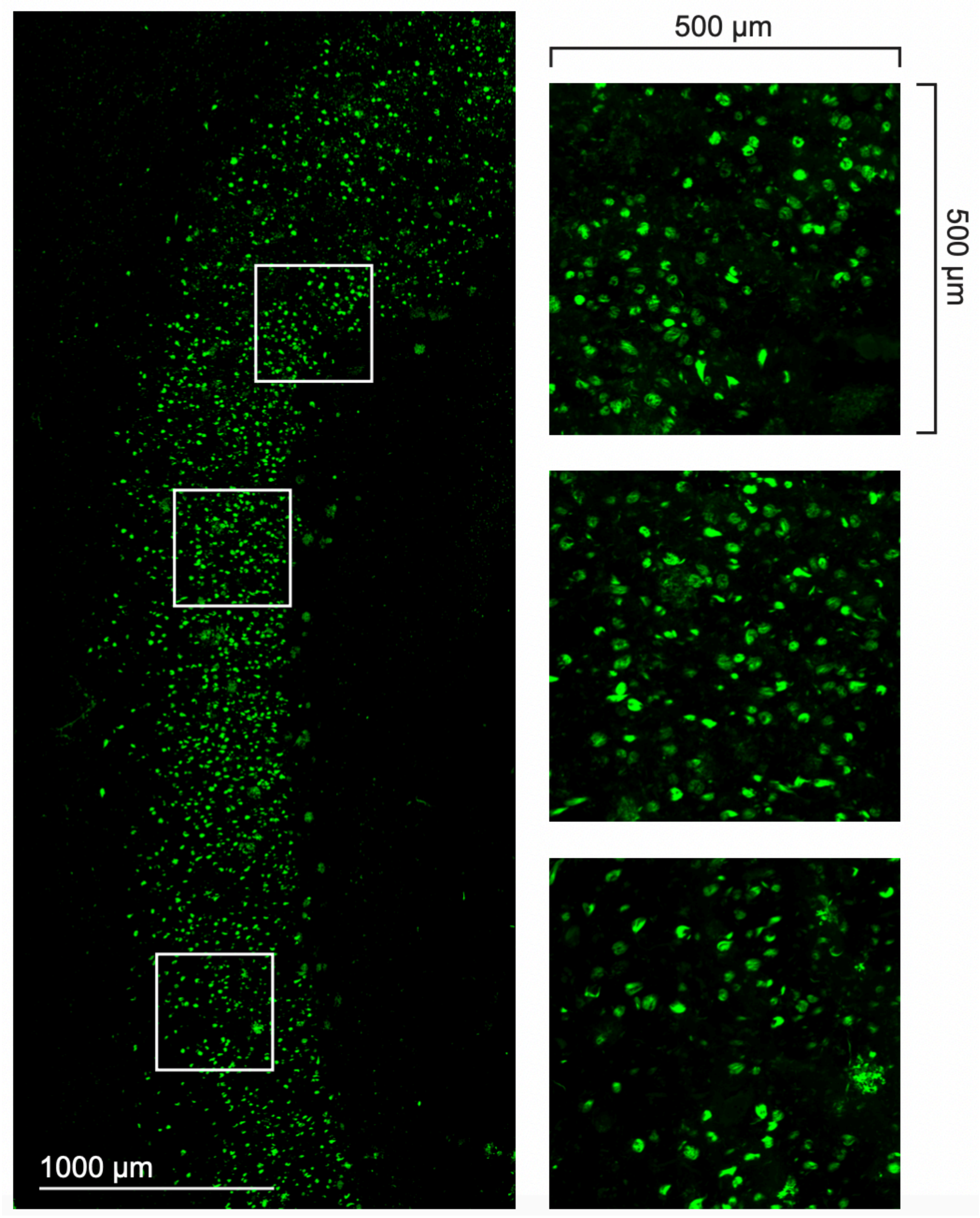

**Supplemental Fig 2** Results of Principal Component Analysis. Principal Component 1 accounted for 59.42% of the variance in regional density, and Principal Component 2 accounted for 18.81% of the variance.

**a.** Variable factor loadings for retained components of regional NFT density principal component analysis. Principal Component 1 may be interpreted as roughly indicating the overall degree of tau pathology accumulation. Principal Component 2 may be interpreted as separating higher hippocampal NFT burden (positive values) relative to higher cortical NFT burden (negative values).

**b.** Individual participant loadings for each principal component by diagnosis. Atypical clinical variants are highlighted. Corticobasal syndrome, logopenic variant primary progressive aphasia, and posterior cortical atrophy can be differentiated from the typical amnestic syndrome and the cognitively normal/very mild dementia group, whereas the individuals with behavioral variant frontotemporal dementia are not clearly discriminable.

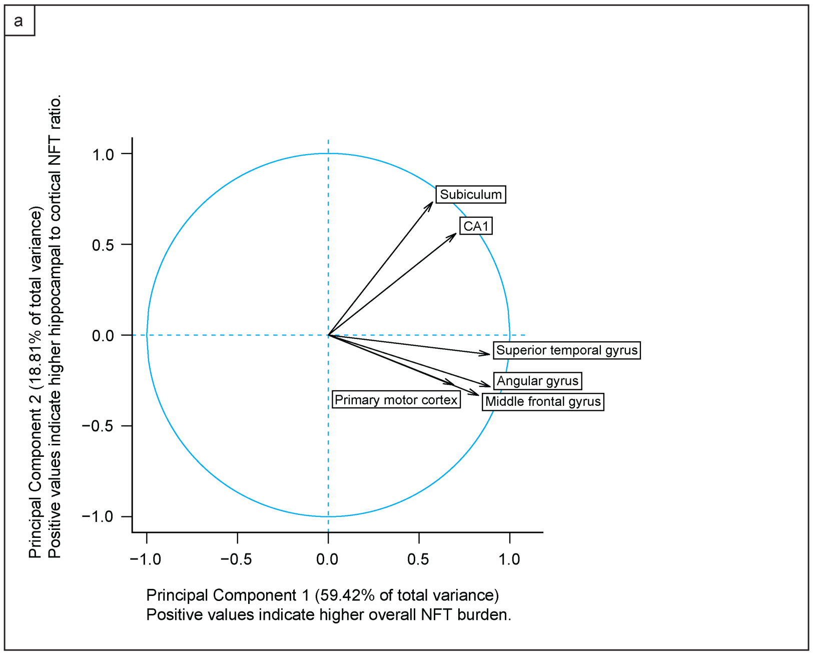


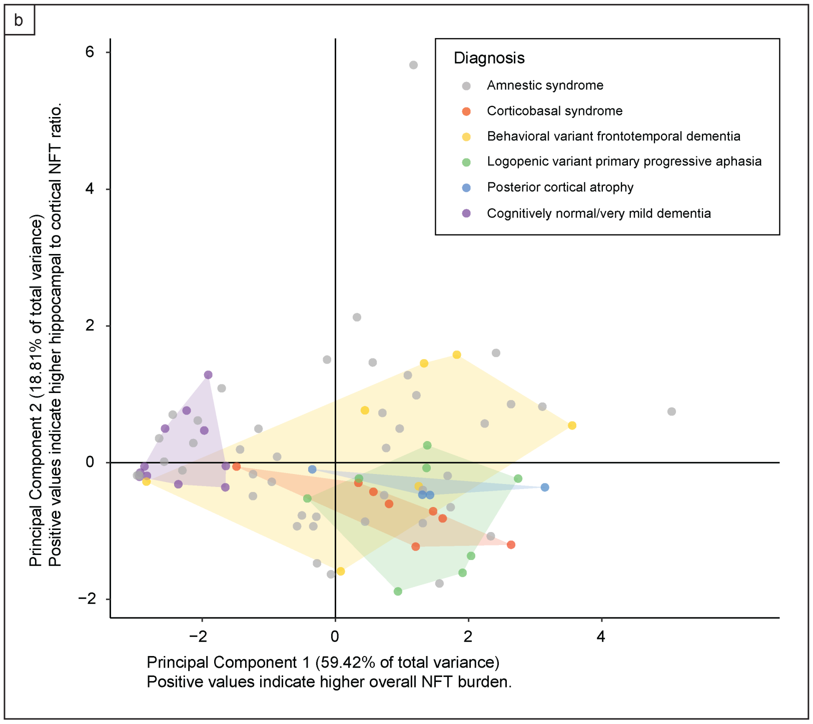
