## Supplemental data for "Alzheimer’s disease clinical variants show distinct regional patterns of neurofibrillary tangle accumulation"

**Journal:** *Acta Neuropathologica*

**Send correspondence to:**

Lea Tenenholz Grinberg, MD, PhD, Associate Professor in Residence

Memory and Aging Center, Weill Institute for Neurosciences, University of California, San Francisco

Data for amnestic cases

| | | | | | | Neurofibrillary tangle density (per mm$^2$) | | | | | |
|---|---|---|---|---|---|---|---|---|---|---|---|
| Case | Sex | Age | Diagnosis | Braak stage | CDR | Middle frontal gyrus | Superior temporal gyrus | Primary motor cortex | Angular gyrus | CA1 | Subiculum |
| 1 | Male | 73 | Amnestic | 6 | 2 | 4.00 | NA | NA | NA | 5.33 | 33.33 |
| 2 | Male | 77 | Amnestic | 6 | 1 | 29.33 | 17.33 | 1.33 | 18.67 | 85.33 | 101.33 |
| 3 | Male | 63 | Amnestic | 6 | 3 | 157.33 | 302.67 | NA | 161.33 | 102.67 | 134.67 |
| 4 | Male | 90 | Amnestic | 6 | 1 | 5.33 | NA | 6.67 | 0.00 | 25.33 | 17.33 |
| 5 | Male | 61 | Amnestic | 6 | 2 | 100.00 | NA | 73.33 | 117.33 | 173.33 | 102.67 |
| 6 | Male | 84 | Amnestic | 6 | 1 | 86.67 | 148.00 | 44.00 | 142.67 | 97.33 | 118.67 |
| 7 | Male | 58 | Amnestic | 6 | 3 | 248.00 | 214.67 | 90.67 | 150.67 | 73.33 | 110.67 |
| 8 | Female | 55 | Amnestic | 6 | 2 | 189.33 | 153.33 | 152.00 | 117.33 | 58.67 | 154.67 |
| 9 | Male | 69 | Amnestic | 6 | 2 | 70.67 | 57.33 | 16.00 | 68.00 | 24.00 | 72.00 |
| 10 | Male | 62 | Amnestic | 6 | 3 | 112.00 | 125.33 | 69.33 | 98.67 | 105.33 | 254.67 |
| 11 | Male | 70 | Amnestic | 6 | 1 | 72.00 | 84.00 | 141.33 | 92.00 | 108.00 | 180.00 |
| 12 | Male | 82 | Amnestic | 4 | 2 | 1.33 | 0.00 | 2.67 | 0.00 | 0.00 | 1.33 |
| 13 | Male | 77 | Amnestic | 6 | 2 | 114.67 | 132.00 | 41.33 | 125.33 | 0.00 | 9.33 |
| 14 | Female | 65 | Amnestic | 6 | 0.5 | 68.00 | 296.00 | 118.67 | 141.33 | 152.00 | 197.33 |
| 15 | Female | 82 | Amnestic | 6 | 2 | 30.67 | 96.00 | 10.67 | 78.67 | 176.00 | 102.67 |
| 16 | Female | 60 | Amnestic | 6 | 1 | 104.00 | 162.67 | 126.67 | 133.33 | 116.00 | 65.33 |
| 17 | Male | 72 | Amnestic | 6 | 1 | 150.67 | 125.33 | 189.33 | 176.00 | 245.33 | 153.33 |
| 18 | Male | 61 | Amnestic | 6 | 1 | 62.67 | 140.00 | 112.00 | 141.33 | 61.33 | 36.00 |
| 19 | Male | 66 | Amnestic | 6 | 1 | 112.00 | 138.67 | 189.33 | 206.67 | 49.33 | 34.67 |
| 20 | Female | 81 | Amnestic | 6 | 2 | 76.00 | 97.33 | 38.67 | 121.33 | 34.67 | 29.33 |
| 21 | Male | 87 | Amnestic | 6 | 3 | 36.00 | 84.00 | 37.33 | 97.33 | 22.67 | 70.67 |
| 22 | Male | 77 | Amnestic | 6 | 0.5 | 85.33 | 130.67 | 74.67 | 129.33 | 141.33 | 109.33 |
| 23 | Male | 56 | Amnestic | 6 | 0.5 | 136.00 | 86.67 | 44.00 | 102.67 | 18.67 | 53.33 |
| 24 | Female | 60 | Amnestic | 6 | 1 | 190.67 | 109.33 | 25.33 | 118.67 | 5.33 | 9.33 |
| 25 | Male | 90 | Amnestic | 6 | 2 | 8.00 | 36.00 | 45.33 | NA | 125.33 | 104.00 |
| 26 | Male | 88 | Amnestic | 6 | 1 | 14.67 | 65.33 | 29.33 | 32.00 | NA | NA |
| 27 | Female | 73 | Amnestic | 6 | 1 | 38.67 | 97.33 | 41.33 | 138.67 | 120.00 | 220.00 |
| 28 | Male | 82 | Amnestic | 6 | 2 | 1.33 | 29.33 | 10.67 | 13.33 | 70.67 | 42.67 |
| 29 | Male | 76 | Amnestic | 6 | 1 | 122.67 | 150.67 | 69.33 | 118.67 | 93.33 | 52.00 |
| 30 | Female | 75 | Amnestic | 6 | 0.5 | 40.00 | 42.67 | 144.00 | 129.33 | 57.33 | 29.33 |
| 31 | Male | 88 | Amnestic | 6 | 2 | 16.00 | 14.67 | 10.67 | 24.00 | 58.67 | 24.00 |
| 32 | Male | 68 | Amnestic | 6 | 1 | 69.33 | 0.00 | 12.00 | 73.33 | 46.67 | 82.67 |
| 33 | Male | 85 | Amnestic | 5 | 1 | 0.00 | 0.00 | 0.00 | 5.33 | 53.33 | 58.67 |
| 34 | Female | 58 | Amnestic | 6 | 1 | 133.33 | 145.33 | 130.67 | 149.33 | 86.67 | 60.00 |
| 35 | Male | 99 | Amnestic | 6 | 1 | 1.33 | 0.00 | 38.67 | 0.00 | 21.33 | 22.67 |
| 36 | Male | 72 | Amnestic | 6 | 0 | 138.67 | 113.33 | 8.00 | 82.67 | 13.33 | 36.00 |
| 37 | Male | 86 | Amnestic | 5 | 1 | 1.33 | 0.00 | 0.00 | 8.00 | 17.33 | 57.33 |
| 38 | Female | 64 | Amnestic | 6 | 2 | 108.00 | 164.00 | 212.00 | 102.67 | 64.00 | 189.33 |
| 39 | Male | 78 | Amnestic | 6 | 2 | 10.67 | 60.00 | 44.00 | 64.00 | 57.33 | 100.00 |
| 40 | Male | 82 | Amnestic | 6 | 2 | 62.67 | 32.00 | NA | 101.33 | 136.00 | 218.67 |
| 41 | Male | 71 | Amnestic | 6 | 2 | 62.67 | 62.67 | 64.00 | 68.00 | 116.00 | 296.00 |
| 42 | Male | 64 | Amnestic | 6 | 3 | 77.33 | 14.67 | 77.33 | 61.33 | 69.33 | 77.33 |
| 43 | Male | 59 | Amnestic | 6 | 3 | 97.33 | 46.67 | 5.33 | 73.33 | 38.67 | 21.33 |
| 44 | Male | 81 | Amnestic | 3 | 1 | 0.00 | 1.33 | 8.00 | 0.00 | 1.33 | 1.33 |
| 45 | Male | 84 | Amnestic | 3 | NA | 0.00 | 1.33 | 2.67 | 1.33 | 2.67 | 0.00 |
| 46 | Female | 78 | Amnestic | 6 | 3 | 76.00 | 165.33 | 25.33 | 162.67 | 112.00 | 204.00 |
| 47 | Female | 67 | Amnestic | 6 | 1 | 130.67 | 193.33 | 32.00 | 170.67 | 180.00 | 252.00 |
| 48 | Female | 68 | Amnestic | 6 | 1 | 5.33 | 30.67 | 8.00 | 37.33 | 9.33 | 32.00 |
| 49 | Female | 64 | Amnestic | 6 | 3 | 306.67 | 221.33 | 100.00 | 266.67 | 196.00 | 316.00 |
| 50 | Female | 66 | Amnestic | 6 | 0.5 | 230.67 | 192.00 | NA | 276.00 | 82.67 | 197.33 |
| 51 | Female | 89 | Amnestic | 6 | 1 | 8.00 | 96.00 | 13.33 | 12.00 | 241.33 | 562.67 |
| 52 | Female | 64 | Amnestic | 6 | 3 | 166.67 | 174.67 | 65.33 | 154.67 | 190.67 | 125.33 |

Data for non-amnestic cases

| | | | | | | Neurofibrillary tangle density (per mm$^2$) | | | | | |
|---|---|---|---|---|---|---|---|---|---|---|---|
| Case | Sex | Age | Diagnosis | Braak stage | CDR | Middle frontal gyrus | Superior temporal gyrus | Primary motor cortex | Angular gyrus | CA1 | Subiculum |
| 53 | Male | 51 | bvFTD | 6 | 2 | 92.00 | 94.67 | 120.00 | 40.00 | 152.00 | 105.33 |
| 54 | Male | 70 | bvFTD | 6 | NA | 101.33 | 158.67 | 20.00 | 125.33 | 165.33 | 193.33 |
| 55 | Male | 69 | bvFTD | 6 | NA | 181.33 | 130.67 | 269.33 | 168.00 | 144.00 | 297.33 |
| 56 | Male | 68 | bvFTD | 6 | 0.5 | 161.33 | 101.33 | 101.33 | 146.67 | 101.33 | 106.67 |
| 57 | Female | 58 | bvFTD | 6 | 3 | 114.67 | 208.00 | 74.67 | 100.00 | 77.33 | 350.67 |
| 58 | Male | 66 | bvFTD | 4 | 2 | 2.67 | 2.67 | 21.33 | 1.33 | 0.00 | 1.33 |
| 59 | Male | 69 | bvFTD | 6 | 3 | 205.33 | 77.33 | 0.00 | 164.00 | 16.00 | 16.00 |
| 60 | Female | 56 | CBS | 6 | 1 | 265.33 | 156.00 | 141.33 | 180.00 | 76.00 | 133.33 |
| 61 | Female | 69 | CBS | 6 | 1 | 174.67 | 166.67 | 105.33 | 152.00 | 64.00 | 113.33 |
| 62 | Male | 64 | CBS | 4 | 2 | 100.00 | 86.67 | 28.00 | 181.33 | 62.67 | 100.00 |
| 63 | Male | 78 | CBS | 6 | 3 | 50.67 | 34.67 | 58.67 | 33.33 | 32.00 | 65.33 |
| 64 | Female | 58 | CBS | 6 | 2 | 80.00 | 141.33 | 244.00 | 124.00 | 52.00 | 141.33 |
| 65 | Female | 61 | CBS | 5 | 0.5 | 96.00 | 138.67 | 180.00 | 165.33 | 37.33 | 85.33 |
| 66 | Female | 79 | CBS | 6 | 0.5 | 52.00 | 178.67 | 141.33 | 96.00 | 29.33 | 126.67 |
| 67 | Male | 78 | CBS | 6 | 1 | 110.67 | 56.00 | 176.00 | 130.67 | 97.33 | 69.33 |
| 68 | Male | 63 | lvPPA | 6 | 3 | 82.67 | 164.00 | 110.67 | 152.00 | 128.00 | 84.00 |
| 69 | Male | 72 | lvPPA | 6 | 2 | 45.33 | 97.33 | 85.33 | 104.00 | 45.33 | 48.00 |
| 70 | Female | 76 | lvPPA | 6 | 3 | 182.67 | 196.00 | 37.33 | 250.67 | 160.00 | 104.00 |
| 71 | Female | 65 | lvPPA | 6 | 2 | 149.33 | 205.33 | 124.00 | 204.00 | 60.00 | 74.67 |
| 72 | Female | 66 | lvPPA | 6 | 2 | 126.67 | 169.33 | 94.67 | 109.33 | 116.00 | 138.67 |
| 73 | Female | 61 | lvPPA | 6 | 2 | 184.00 | 194.67 | 173.33 | 130.67 | 88.00 | 12.00 |
| 74 | Male | 58 | lvPPA | 6 | 2 | 136.00 | 176.00 | 130.67 | 148.00 | 17.33 | 14.67 |
| 75 | Female | 72 | lvPPA | 6 | 2 | 148.00 | 161.33 | 38.67 | 60.00 | 86.67 | 61.33 |
| 76 | Male | 66 | Other | 6 | 3 | 16.00 | 25.33 | 9.33 | 18.67 | 26.67 | 32.00 |
| 77 | Female | 58 | PCA | 6 | 1 | 136.00 | 172.00 | 89.33 | 153.33 | 80.00 | 110.67 |
| 78 | Female | 63 | PCA | 6 | 3 | 128.00 | 126.67 | 94.67 | NA | 120.00 | 32.00 |
| 79 | Female | 65 | PCA | 6 | 1 | 60.00 | 50.67 | 126.67 | 81.33 | 45.33 | 116.00 |
| 80 | Male | 59 | PCA | 6 | 2 | 158.67 | 101.33 | NA | 138.67 | 58.67 | 64.00 |
| 81 | Female | 67 | PCA | 6 | 3 | 257.33 | 101.33 | 156.00 | 193.33 | 232.00 | 58.67 |
| 82 | Female | 68 | PCA | 6 | 3 | 92.00 | 145.33 | 32.00 | 108.00 | 66.67 | NA |
| 83 | Female | 62 | PCA | 6 | 3 | 154.67 | 121.33 | 128.00 | 138.67 | 56.00 | 154.67 |
| 84 | Female | 81 | VMD | 2 | 0.5 | 13.33 | 8.00 | 1.33 | 0.00 | 2.67 | 5.33 |
| 85 | Female | 86 | VMD | 3 | 0.5 | 1.33 | 0.00 | 6.67 | 1.33 | 1.33 | 0.00 |
| 86 | Female | 89 | VMD | 3 | 0.5 | 17.33 | 5.33 | 42.67 | 13.33 | 57.33 | 61.33 |
| 87 | Male | 76 | VMD | 2 | 0.5 | 9.33 | 10.67 | 58.67 | 8.00 | 20.00 | 1.33 |
| 88 | Female | 87 | VMD | 4 | 0.5 | 5.33 | 4.00 | 8.00 | 9.33 | 60.00 | 69.33 |
| 89 | Male | 78 | VMD | 6 | 0.5 | 8.00 | 6.67 | 5.33 | 13.33 | 76.00 | 124.00 |
| 90 | Male | 81 | VMD | 5 | 0.5 | 34.67 | 44.00 | 22.67 | 48.00 | 46.67 | 29.33 |
| 91 | Male | 76 | CN | 2 | 0 | 2.67 | 2.67 | 1.33 | 0.00 | 4.00 | 2.67 |
| 92 | Male | 77 | CN | 5 | 0 | 2.67 | 0.00 | 8.00 | 0.00 | 1.33 | 21.33 |
| 93 | Male | 103 | CN | 4 | 0 | 5.33 | 0.00 | 0.00 | 1.33 | 40.00 | 48.00 |
| 94 | Male | 91 | CN | 4 | 0 | 44.00 | 40.00 | 18.67 | 65.33 | 29.33 | 20.00 |
